# Fusion of Drosophila oocytes with specified germline sister cells

**DOI:** 10.1101/2020.01.22.915736

**Authors:** Zehra Ali-Murthy, Richard D. Fetter, Thomas B. Kornberg

**Affiliations:** Cardiovascular Research Institute, University of California, San Francisco, CA 94143; Department of Biochemistry and Biophysics, University of California, San Francisco, CA 94143

**Keywords:** Drosophila, oocyte, nurse cell, cell fusion

## Abstract

In many animals, oocytes develop together with sister germline cells that pass products to the developing oocyte. In Drosophila, fifteen sister germline (nurse) cells in each egg chamber are known to apoptose by stage 12-13, but we discovered that two specific nurse cells that are juxtaposed to the oocyte are eliminated precociously at stage 10B. These nurse cells fuse with the oocyte and their nuclei extrude through an opening that forms in the oocyte. These nuclei extinguish in the ooplasm, and at stage 11, egg chambers have thirteen nucleated nurse cells and the plasma membrane of the oocyte is mostly restored. In infrequent egg chambers in which nurse cells are not eliminated, oocytes do not develop normally and are not fertilized. Precocious elimination is common to other Drosophila species. We conclude that nurse cells are distinguished by position and identity, and that nurse cell dissolution proceeds in two stages.

## Introduction

The eggs of many species are distinguished by specialized coatings, structures and organelles that enable fertilization and early development. Another distinctive feature is the provenance of egg components, of which some are produced by the oocyte and some by other cells. The external sources in nearly all egg-laying vertebrates and invertebrates include somatic cells such as those that contribute yolk proteins. The external sources also include germline cells whose destiny is not fertilization and zygote formation. Instead, these cells export products they make to provision a developing oocyte, and die rather than mature to fertilization competence, despite their germline lineage. Hydra is an example with such sacrificial germline cells; early Hydra oocytes fuse with nearby germ cells that differentiate as nurse cells and transfer cytoplasm directly to the growing oocyte (Alexandrova et al., 2005; Miller et al., 2000). Another example is the mouse oocyte that receives cytoplasmic components including organelles such as centrioles, Golgi, and mitochondria from neighboring, interconnected sister germline cells. These nurse-like cells are eliminated by an apoptososis-like process (Lei and Spradling, 2016).

Drosophila oogenesis is arguably the best understood example. Four consecutive mitotic divisions of a primary oogonial cell generate a cyst of sixteen cells. One of the daughters of the first division develops as the oocyte while the other fifteen descendants of the primary oogonial cell develop into nurse cells. These sixteen clonally-related germline cells have stable intercellular bridges (“ring canals”) that form at sites of arrested cleavage furrows, linking sister cells (Matova and Cooley, 2001a). Although both the oocyte and nurse cells and their nuclei increase in volume as oogenesis proceeds, the respective developmental programs for the two cell types differ. The nurse cells undergo 10-12 endocycles without cytokinesis, becoming increasingly polyploid between stages 2-10 (for descriptive purposes, oogenesis is categorized by 14 morphologically-distinct stages) (Dej and Spradling, 1999). In contrast, the chromosomes of the oocyte nucleus (the “germinal vesicle”) proceed through meiosis, and although the germinal vesicle increases in size, it does not endoreplicate. The ooplasm is understood be constituted mostly of nurse cell products that the nurse cells export to the oocyte through the ring canals, a process termed “dumping”. Although the more posterior nurse cell nuclei have higher ploidy and have larger nuclei than the more anterior nurse cells, there has been no evidence to suggest that different nurse cells have distinct roles or fates.

The first overt signs of incipient nurse cell death have been noted in stage 10B, when each nurse cell nucleus becomes surrounded by a dense actin network, the nuclear membrane transforms from a smooth and spherical shape to an irregular morphology with multiple folds, and the permeability of the nuclear membrane increases (Cooley et al., 1992). These changes apparently facilitate and are necessary for rapid transport of nurse cell cytoplasm and nucleoplasm through the ring canals to the oocyte (Guild et al., 1997; Mahajan-Miklos and Cooley, 1994). Subsequently, nurse cell nuclei condense and degrade, and the remnants are cleared, leaving only the oocyte in the egg chamber at stage 14. Because genetic studies have not implicated the canonical inducers and inhibitors of apoptosis in the nurse cell death program (Baum et al., 2007; Foley and Cooley, 1998; Peterson et al., 2003), the mechanism of killing has not been categorized (Peterson and McCall, 2013).

This study describes a new feature of the nurse cell program. We discovered that nurse cell nuclei are absorbed into the oocyte at late stage 10B, prior to dumping and prior the phase of nuclear condensation. This process involves reorganization of the oocyte plasma membrane, membrane fusion with nurse cells, and long range movement of nurse cell nuclei into the ooplasm. These nurse cell nuclei degrade prior to stage 11. Because the identities of these nurse cells are defined, we infer that different nurse cells have distinct identities and roles. We present evidence that this unprecedented type of cell fusion and cell elimination program is essential for oocyte maturation, and speculate on its purpose.

## Results

### Nurse cell nuclei enter the late stage 10B oocyte

The adult ovary is an assembly of approximately 20 production lines (ovarioles) that each contain a string of egg chambers. Every egg chamber has 16 germline descendants of a single oogonial cell, and is encapsulated by a layer of somatic follicle cells. Classification of 14 stages of oogenesis that are represented in these egg chambers is based on morphology, and is not related to duration - the stages vary from 0.5 to 6 hours (Dej and Spradling, 1999). And because the number of egg chambers per ovariole averages less than nine, not all stages are present in every ovariole or in every ovary. Checkpoints that control oocyte maturation influence the distribution of oocyte stages (Drummond-Barbosa and Spradling, 2001) and lead to over- or under-representation of particular stages in individual ovaries (Fig. 1A). Although late stage 10B oocytes are rare in females raised under standard conditions of fly culture (∼0.4/ovary), we developed a regimen to increase their frequency approximately 2.8 times to an average of 1.1 per ovary (Fig. 1B).

**Figure 1.**
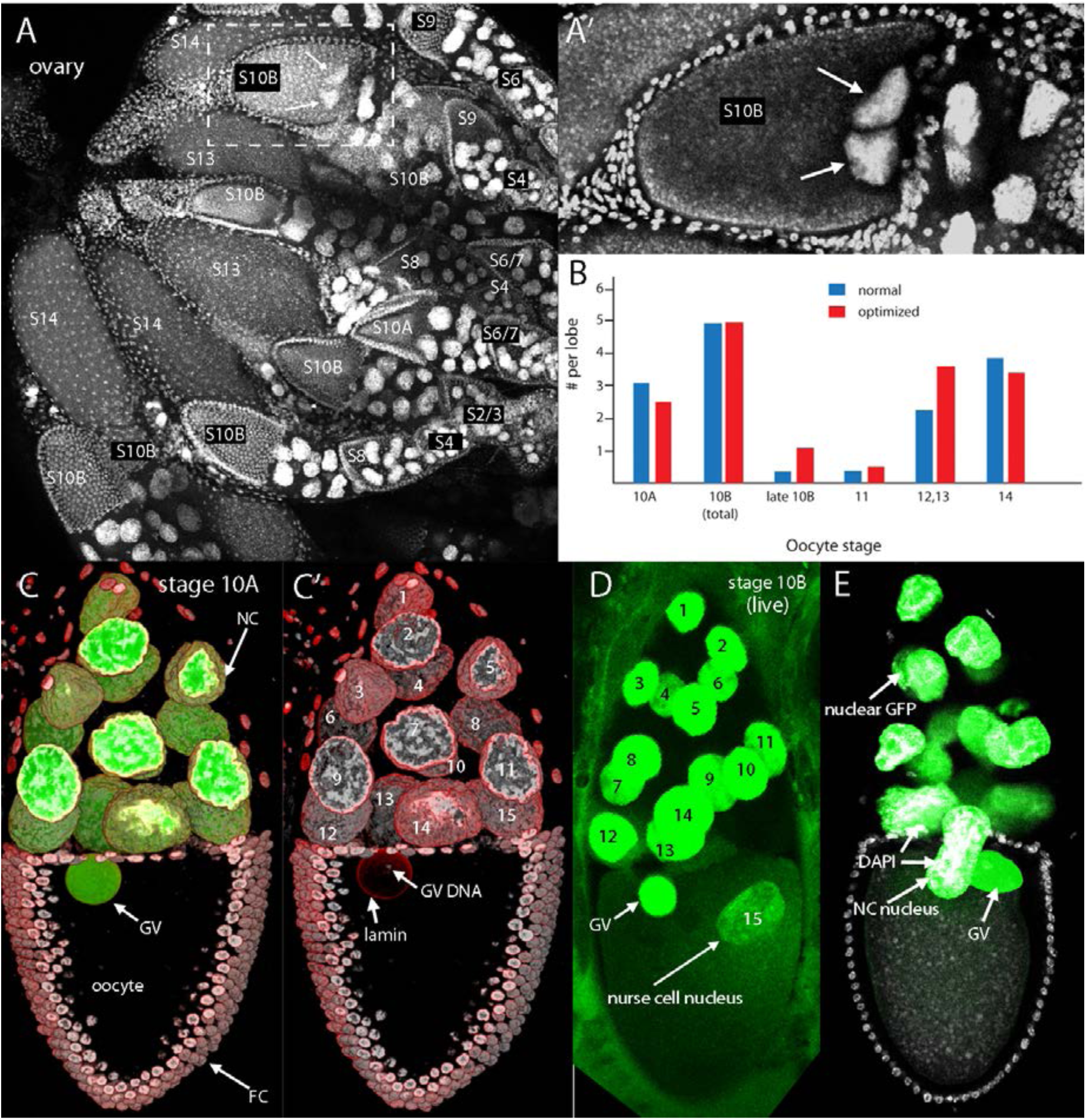
Nurse cell nuclei in stage 10 oocytes. (A) WT ovary stained with DAPI, egg chamber stages indicated; small follicle cell nuclei cover oocyte that is adjacent to region with large polyploid nurse cell nuclei in each egg chamber. (A’) higher magnification image of stage 10B egg chamber (boxed by white-dashed lines in (A)), arrows indicate nurse cell nuclei in ooplasm. (B) graph tabulating egg chamber stages in WT ovaries of females raised under normal conditions or conditions optimized for stages late 10B (with nurse cell nuclei in ooplasm) and 11. (C,C’) Stage 10A egg chambers with germline expression of nuclear GFP (green) in 15 nurse cell nuclei and germinal vesicle (GV), stained with ⍰-lamin antibody (red) and DAPI (white); nurse cell (NC). (D) Unfixed (live) stage 10B egg chamber with germline expression of nuclear GFP (green) in 15 nurse cell nuclei, one in ooplasm (arrow). (E) Fixed stage 10B egg chamber stained with DAPI (white) with germline expression of nuclear GFP, and one entering nurse cell nucleus (arrow).

The nucleus (germinal vesicle, GV) of stage 10A oocytes localizes near to the anterior face of the oocyte. The GV and nurse cell nuclei can be visualized by staining with anti-nuclear lamin antibody and by fluorescence of GFP in nuclei of egg chambers dissected from ovaries of flies with germline expression of nuclear GFP (Fig. 1C,C’). The GV and nurse cell nuclei are similar in size, but whereas most of the volume of a nurse cell nucleus stains with DAPI, DAPI staining DNA occupies only a small fraction of the GV. In many late stage 10B egg chambers, oval-shaped and elongated fluorescent nuclei are also visible in the oocyte, in addition to the nuclear GFP fluorescence of the GV. The egg chamber in Figure 1D that has one such “ectopic” nucleus was imaged without fixation. Imaging late stage 10B egg chambers after fixation and DAPI staining reveals that these structures have DNA content that is typical of nurse cells (Fig. 1E). The whole ovary micrograph in Figure 1A has a stage 10B oocyte with two of these DAPI-positive structures (Fig. 1A,A’). Evidence described in the following sections shows that these structures are nurse cell nuclei that migrate into the oocyte.

Because the mitoses that generate the 16 germline cells of the egg chamber are stereotyped and characterized by incomplete cytokinesis, the lineage of each cell is unique and each cell is distinguished by the number of ring canals that link it to its sisters. For example, both cells produced by the first division have four ring canals, and the first daughters of these two cells each have three (Fig. 2A, Fig. 2supp). We determined the identities of the 16 germline cells in 24 egg chambers by marking the nuclei with nuclear GFP and staining the ring canals with α-Hts antibody. Two features are noteworthy. First, although the position of each cell is not precisely the same in every egg chamber, the different cells are reproducibly positioned into one of three tiers (Fig. 2 B-M). The four cells that connect directly to the oocyte (designated “tier 1”) are numbers 2, 3, 5, 9. Six cells (numbers 4, 6, 7, 10, 11, 13) are intermediate (“tier 2”), and five cells (numbers 8, 12, 14, 15, 16) are in the region most distant from the oocyte (“tier 3”), and these eleven cells are not in direct contact with the oocyte. These results are consistent with those reported in Alsous et al, 2017 (Imran Alsous et al., 2017). Second, whereas all stage 3-10A egg chambers have 15 nurse cell nuclei, some late stage 10B oocytes have a nurse cell nucleus in the anterior ooplasm, and many late stage 10B egg chambers have only 13 nucleated nurse observed (Fig. 3D,E), but these are rare.

**Figure 2.**
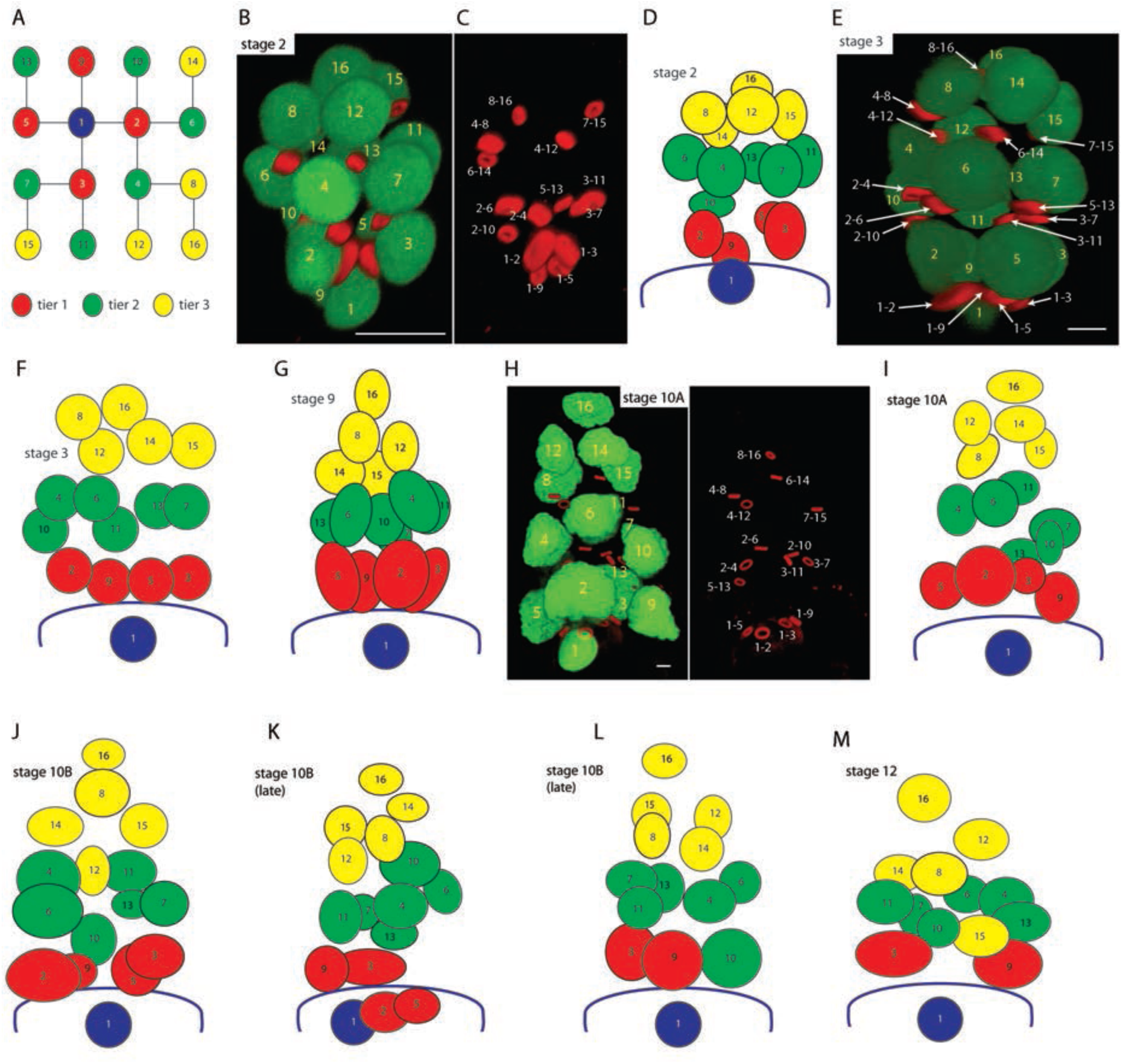
Defined positions of nurse cell nuclei. (A) Lineage tree of nurse cells; oocyte (1; blue), oocyte daughters (2, 3, 5, 9; tier 1, red), oocyte granddaughters (4, 6, 7, 10, 11, 13; tier 2, green), oocyte great granddaughters (8, 12, 14, 15, 16; tier 3, yellow); tiers 1, 2, and 3 are touching the oocyte, in middle nurse cell region, farthest from oocyte, respectively. (B) Stage 2 egg chamber with nuclear GFP expression (green) and ring canals stained with α-HTS antibody, numbered nuclei correspond to nomenclature in (A). (C) Egg chamber in (B) showing α-HTS antibody-stained ring canals with identities of nurse cells each ring canal link. (D) Cartoon of egg chamber in (B). (E) Stage 3 egg chamber with nuclei and ring canals identities marked. (F) Cartoon of egg chamber in (E). (G) Cartoon of stage 9 egg chamber with identities of nuclei marked. (H,I) Stage 10A egg chamber with identities of nuclei and ring canals marked (H) and depicted in cartoon (I). (J-M) Cartoons depicting identities of nuclei in stage 10B - stage 12 egg chambers.

**Figure 3.**
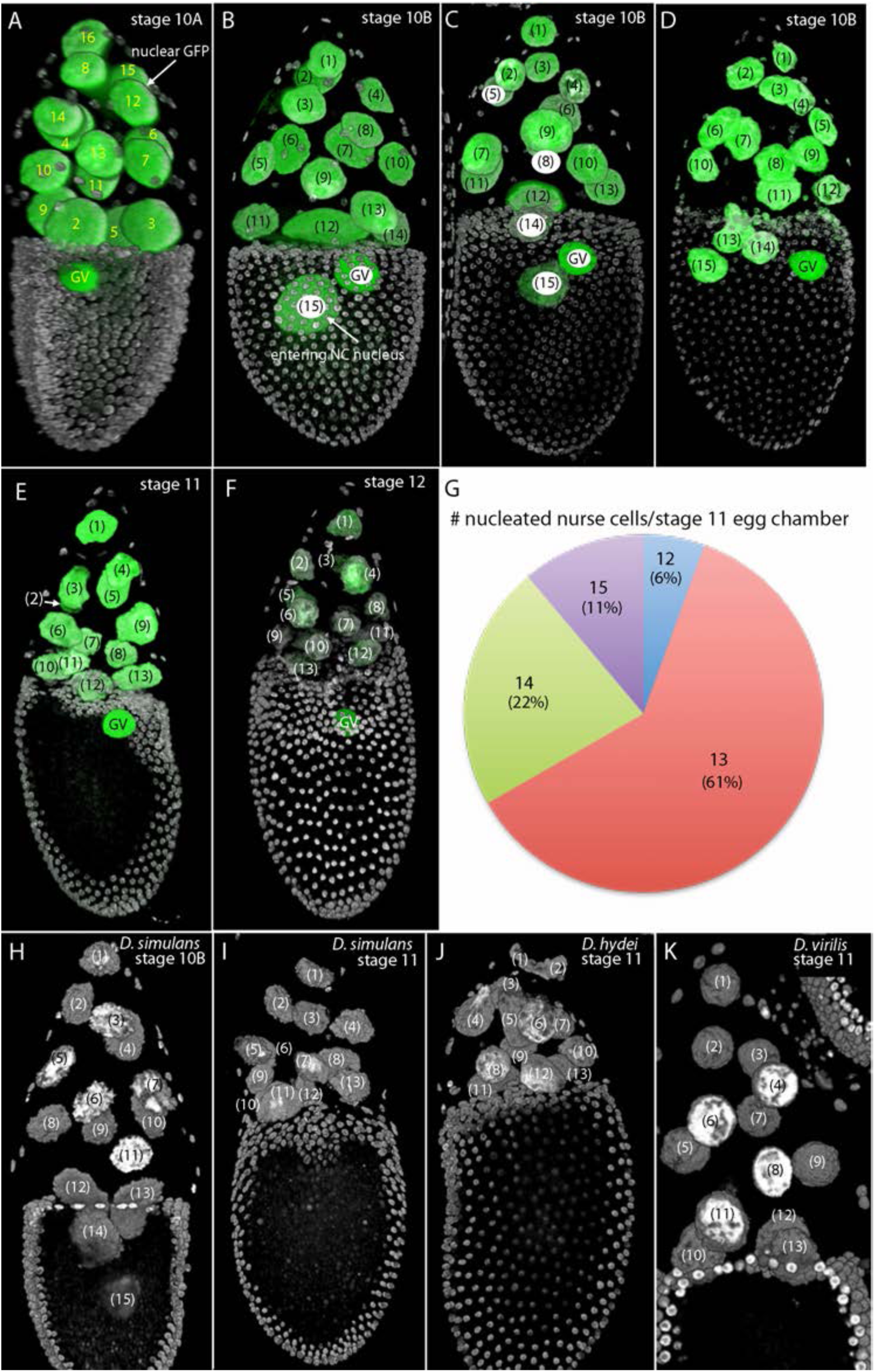
Nurse cell content in Drosophila egg chambers. (A-F) *D. melangaster* stage 10A – stage 12 egg chambers expressing nuclear GFP (green) and stained with DAPI (white), and nuclei identified (A) or counted (numbers in parentheses, B-F); stage 10B egg chambers with one (B), two (C), or three (D) entering nurse cell nuclei, and stage 11 (E) and stage 12 (F) egg chambers with a total of thirteen nurse cell nuclei. (G) Pie chart indicating proportion of stage 11 egg chambers with 12, 13, 14 or 15 nurse cell nuclei. (H-K) *D. simulans*, *D. hydei*, and *D. virilis* egg chambers stained with DAPI (white) with counts of nurse cell nuclei showing 15 nurse cell nuclei with two fully entered into the stage 10B *D. simulans* oocyte (H), and 13 nurse cell nuclei in the *D. simulans* (I), *D. hydei* (J), and *D. virilis* (K) stage 11 egg chambers.

This description is based on our analysis of ovaries that were fixed immediately following removal of ovaries from the female and prior to dissection of the ovary. Following fixation, ovary dissection involved teasing ovarioles apart with fine needles in ways that minimized effects on the shape or structure of the egg chambers. In order to substantiate the unexpected redistribution of nurse cell nuclei in stage 10B egg chambers, we analyzed eighteen stage 11 oocytes in such preparations (Fig. 3G). Stage 11 egg chambers are infrequent, averaging less than one per two ovaries, but among the eighteen ovaries, none had nurse cell nuclei in the ooplasm, eleven (61%) had thirteen nucleated nurse cells, and only two egg chambers (11%) had fifteen nucleated nurse cells. This establishes that the presence of nurse cell nuclei in the ooplasm of stage 10B oocytes is not an artifact of the procedure we used to image the ovarioles.

The reduction in number of nurse cell nuclei is not unique to a particular fly stock or line, but is characteristic of every wild type and mutant *D. melanogaster* stock we examined. It is also a feature of *D. simulans, D. hydei, and D. virilis* wild types (Fig. 3H-K). These images are consistent with the idea that prior to the onset of the program that clears nurse cells from stage 12-14 egg chambers (Foley and Cooley, 1998), a common feature of Drosophila oogenesis involves the absorption of nurse cell nuclei into the ooplasm of late stage 10B oocytes. For the following discussion, we distinguish between egg chambers with fifteen nucleated nurse cells and egg chambers with fewer than fifteen and focus on the majority, which have thirteen.

To determine if the nuclei that enter the oocyte are a random selection from the fifteen nurse cells or are from particular nurse cells, we identified the nucleated nurse cells in both late stage 10B oocytes that have entering nuclei, and in stage 11 egg chambers (Fig. 4A,B). In all the stage 10B egg chambers we examined, the entering nuclei were from nurse cells 2 (the sister of the first mitotic division) and 5 (the sister of the third mitotic division) (Figs. 2K, 4A,B). Both cells 2 and 5 are tier 1 cells, and are directly linked to the oocyte by ring canals. This finding indicates that the process of nuclear migration is specific to particular nurse cells and it is consistent with the idea that at least two of the nurse cells among the fifteen in each egg chamber may be different from the others.

**Figure 4.**
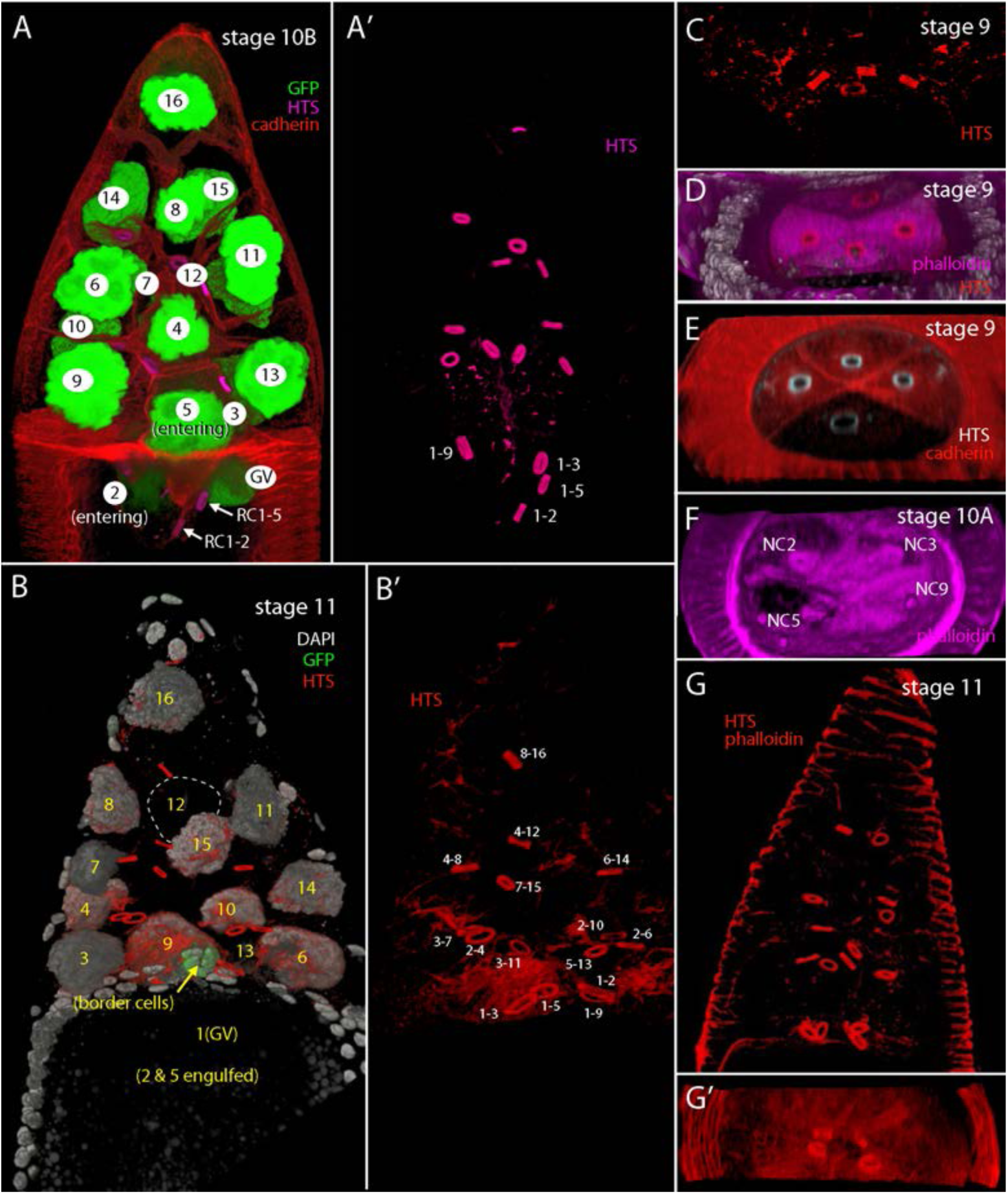
Ring canals precede nurse cell nuclei into oocyte. (A,A’) Stage 10B egg chamber expressing nuclear GFP (green), stained with α-HTS antibody to mark ring canals and α-cadherin antibody to mark cell boundaries, with nuclei identified; ring canals linking oocyte with nurse cell 2 (RC1-2) and nurse cell 5 (RC1-5) are positioned in the oocyte posterior to the entering nuclei of nurse cells 2 and 5. (B,B’) Stage 11 egg chamber lacking absent nurse cell 2 and 5 nuclei, expressing nuclear GFP (green) and stained with DAPI (white) and α-HTS antibody to mark ring canals that are all anterior to the oocyte. (C-F) Transverse optical sections at the anterior face of the oocyte (marked with phalloidin fluorescence (D,F), or α-cadherin antibody (E)) created from 3D projection of confocal optical sections show arrangements of ring canals (marked with α-HTS antibody) and gaps in fluorescence-marked anterior oocyte face at sites of nurse cells 2 and 5 (F). (G, G’) All ring canals anterior to the oocyte and four ring canals at the anterior oocyte face in stage 11 egg chamber marked with α-HTS antibody staining and phalloidin fluorescence (red).

The four tier 1 nurse cells are direct descendants of the oocyte and each tier 1 cell connects directly to the oocyte through a ring canal. At stage 9, the four ring canals are evenly spaced on the anterior face of the oocyte, each ring canal within quadrant of a planar, circular structure that can be imaged by cadherin and phalloidin staining (Fig. 4C-E). This organized structure is temporary. Whereas it stains uniformly across the nurse-cell oocyte interface at stage 9, staining is not continuous at stage 10: gaps are present at nurse cells 2 and 5 (Fig. 4F), and the ring canals that join the oocyte to nurse cells 2 and 5 (RC1-2 and RC1-5) are posterior to the plane of the oocyte anterior face (Fig. 4A).

These ring canals are in the interior of the oocyte, and appear to precede the nearby entering nurse cells 2 and 5. The ring canals are small relative to entering nurse cell nuclei (∼14 μm vs ∼44 μm diameter), and because the fluorescent images are not consistent with a direct association between the nurse cell nuclei and ring canals, we conclude that the nurse cell nuclei do not extrude through the ring canals. At stage 11, RC1-2 and RC1-5 are again at the anterior planar interface of the oocyte, and are again organized in an evenly spaced pattern (Fig. 4B,B’,G,G’).

In these preparations, the fluorescence of nuclear GFP reveals the location of nurse cell nuclei numbers 2 and 5, but the images do not distinguish whether the nuclei are expelled from the nurse cells or if the nurse cells are engulfed by the oocyte in their entirety. We do not have a way to independently mark nurse cells 2 and 5, but three observations are consistent with the idea that nurse cells 2 and 5 remain in their original location. First, whereas ring canals RC1-2 and RC1-5 (which connect nurse cells 2 and 5 with the oocyte) move into the oocyte together with the entering nurse cell nuclei, ring canals RC2-4, RC2-6, RC2-10 and RC5-13 do not appear to change their relative location (Fig. 4A,A’, Fig. 5A,A’). This is consistent with the idea that the morphology of nurse cells 2 and 5 at the juxtapositions with nurse cells 4, 6, 10, and 13 do not change. Second, the process of nuclear extrusion appears to leave nurse cells 2 and 5 as enucleated structures, and the adjoining cells do not expand into the spaces that the nuclei of nurse cells 2 and 5 vacated at stage 10B. These spaces are devoid of nuclei in stage 11 egg chambers (Fig. 5B-D’). Third, the nuclei of nurse cells 2 and 5 are large and their movement into the oocyte might be expected to increase the volume of the oocyte when they enter. We measured the volumes of egg chambers and oocytes from stage 2-13, and determined that the egg chamber volume and oocyte volume increase with doubling times of 6.7 hours and 5.5 hours, respectively (Fig. 5E). As noted in previous studies, oocyte volume increases more rapidly during the stage 11 to stage 14 period of “dumping” when the contents of the nurse cells become subsumed into the oocyte (Guild et al., 1997). Our measurements indicate that the oocyte doubling time is 5.9 hours between stages 2 to 11 and 2.8 hours for stages 11-13. The growth rate of the oocyte did not change significantly at stage 10B when the nuclei of nurse cells 2 and 5 enter the oocyte, consistent with the idea that the nuclei of nurse cells 2 and 5 enter the oocyte without changing the cellular composition or arrangement of the nurse cell cluster.

**Figure 5.**
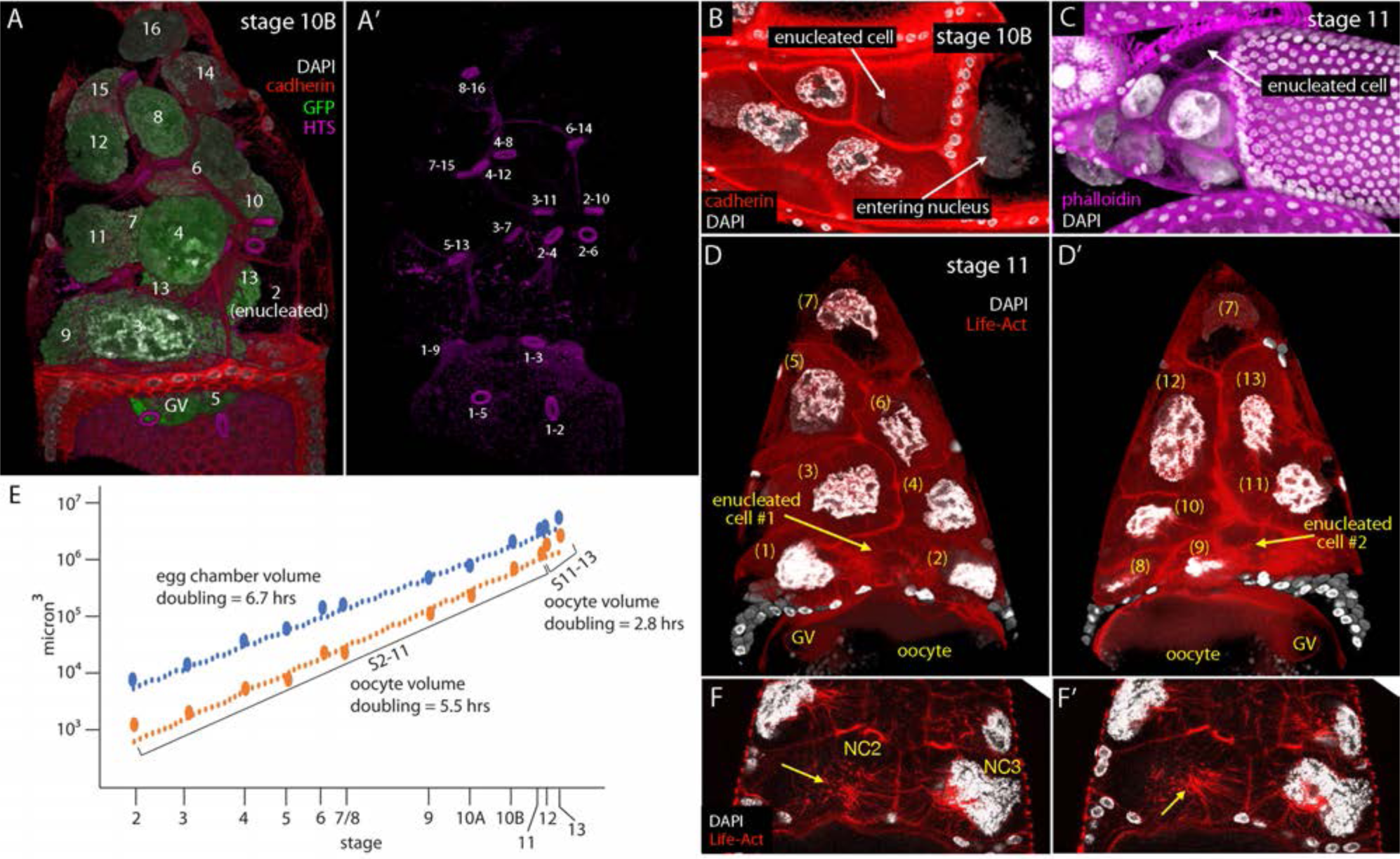
Enucleated nurse cells in stage 10B and stage 11 egg chambers. (A,A’) Stage 10B egg chamber marked with nuclear GFP (green), DAPI (white), α-HTS (purple), and α-cadherin (red), with identified nurse cell nuclei and ring canals, and enucleated nurse cell 2. (B-D) Enucleated nurse cells indicated (arrows) in a stage 10B (B) two stage 11 (C,D,D’) egg chambers. (E) Graph showing volumes of stage 2 – stage 13 egg chambers. (F,F’) Stage 11 egg chamber marked with Life-Act (red) and DAPI (white) fluorescence and arrows indicating actin cables in enucleated nurse cell 2.

Stage 11 nurse cells have actin networks that form “cages” around the nuclei and that appear to attach to the nuclear envelopes. It has been proposed that the actin cages stabilize the nuclei and prevent them from physically blocking the intracellular bridges during dumping (Guild et al., 1997). We observed that enucleated nurse cells have networks of actin cables that are similar to the networks in nucleated nurse cells (Fig. 5F,F’), indicating that formation of the networks is not dependent on nuclei or on the presence of a nuclear envelope.

### A channel connects the oocyte and nurse cells at stage 10B

We investigated whether the nurse cell nuclei might enter the late stage 10B oocyte through an opening in its anterior face. We used various markers to examine the distribution of nurse cell nuclei, ring canals and germinal vesicle with respect to the membranes of follicle cells, border cells, and oocyte (Fig. 6A-F). In the four egg chambers shown in these panels, phalloidin and cadherin staining and fluorescence of membrane-tethered GFP delineated the plasma membranes of the follicle cells and oocyte except in the immediate region of the entering nurse cell nucleus. The apparent opening between the oocyte and follicle cell region is also revealed by the absence of Dpp:Cherry fluorescence that is prominent in the centripetal cells (Fig. 6E,F). The proximity and relative positions of the germinal vesicle (Fig. 6A-C) and border cells (Fig. 6D,F) in these egg chambers are typical.

**Figure 6.**
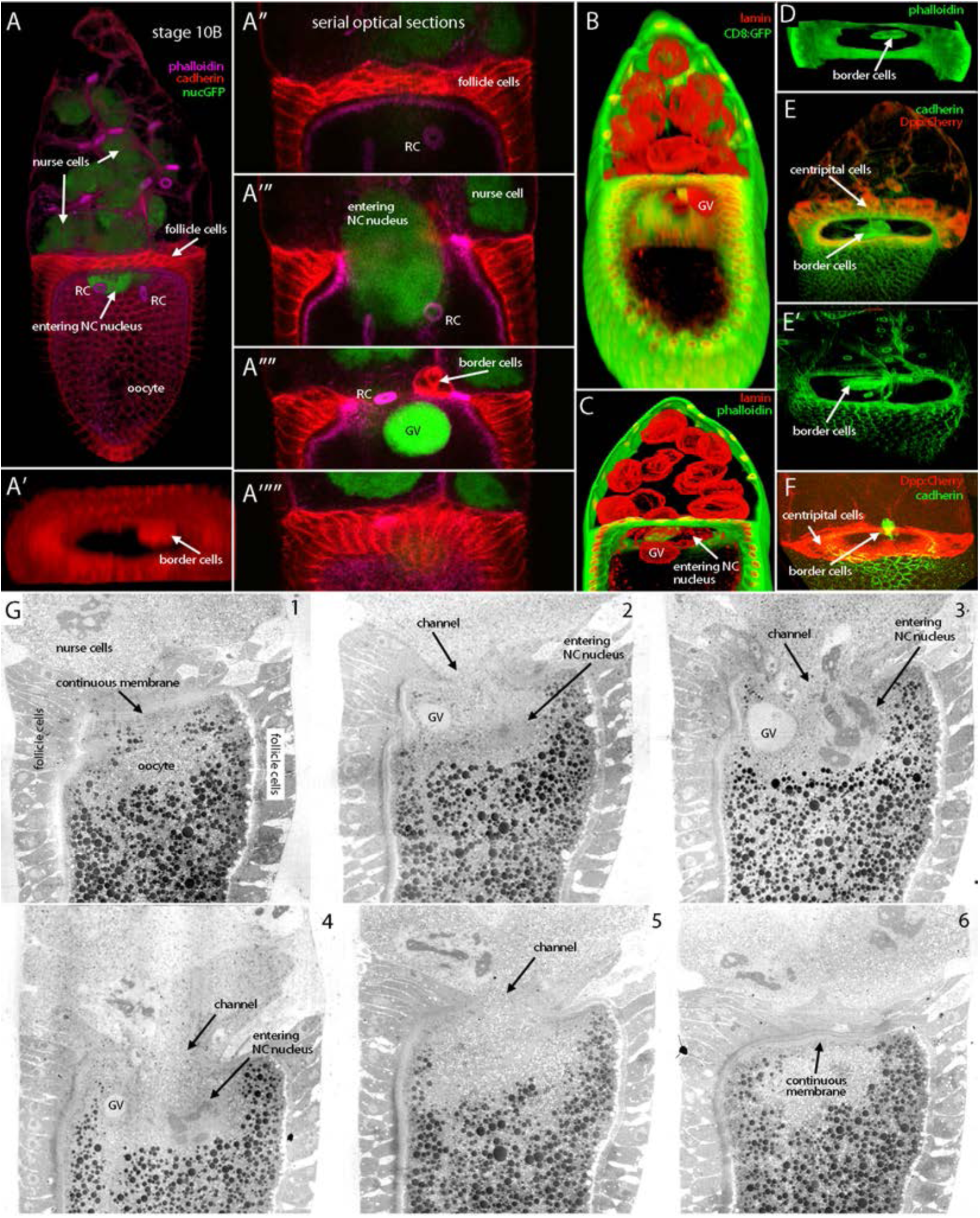
Anatomy of the anterior face of the stage 10B oocyte. (A-A’””) Stage 10B egg chamber marked with nuclear GFP (green), phalloidin fluorescence (purple), and α-cadherin staining identifies oocyte, nurse cells, nurse cell nuclei, follicle cells, border cells and ring canals, but phalloidin or cadherin staining is absent at the anterior face of the oocyte at the site of entering nurse cell nucleus. (B-F) Stage 10B egg chambers with pore at oocyte anterior face, marked (B) with membrane-tethered GFP (green) and α-lamin (red), or (C) α-lamin (red) and phalloidin fluorescence, (D) phalloidin fluorescence, and (E,E’,F) Dpp:Cherry (red) and α-cadherin (green). (G) Electron microscope images of successive sections of a stage 10B oocyte showing continuous plasma membrane across the anterior oocyte face (1, 6) and channel through which a nurse cell nucleus enters (2-5).

These images are examples of our attempts to detect cell plasma membranes in the region where the nurse cell nuclei enter the oocyte. The lack of evidence in these images contrasts with similar images of stage 9 and 11 egg chambers that show phalloidin and cadherin over the anterior oocyte face (Figs. 4, 5). Although these results are consistent with the idea that plasma membranes do not separate the oocyte and nurse cells in this region of late stage 10B egg chambers, we cannot rule out the possibility that confocal fluorescence imaging has insufficient sensitivity to detect membranes in this region. We therefore obtained high resolution images by electron microscopy.

The six sections in Figure 6G show a stage 10B egg chamber at 10 μm intervals. Whereas the continuity of the membrane across the anterior face of the oocyte is evident in sections 1 and 6, the intermediate sections 2-5 do not have plasma membrane in the region of the entering nurse cell nucleus. Instead, the central region of the anterior end of the oocyte appears to directly juxtapose ooplasm and nurse cell cytoplasm. A nurse cell nucleus spans this apparent channel.

To analyze the central region in greater detail, sections of stage 10B egg chambers with entering nurse cell nuclei were imaged at high magnification and montages were assembled that span the region from one side of the oocyte to the other (Fig. 7), or from the entering nurse cell nucleus at the oocyte anterior to the oocyte posterior pole (Fig. 8). The 10 μm interval sections in Figure 7A-F show the arrangement of the border cells, GV, and channel to two entering nurse cell nuclei. The entering nurse cell nucleus on the right side is almost entirely inside the oocyte (Fig. 7C-E); only a small portion of the nurse cell nucleus on the left side is inside (Fig. 7D,E). The germinal vesicle (GV) and border cells are situated in the region where the oocyte and nurse cells are joined (Fig. 7B), but the plasma membranes of the juxtaposed oocyte and nurse cells are not present in this middle region. The montage of high magnification images in Figure 7G shows a complex assembly of interdigitated membrane fingers/protrusions on the left side and no membranes on the right side. The morphology of this egg chamber is therefore consistent with the idea that the ooplasm of the oocyte and cytoplasm of the nurse cells is continuous and that the plasma membranes in the region of juxtaposition have rearranged.

**Figure 7.**
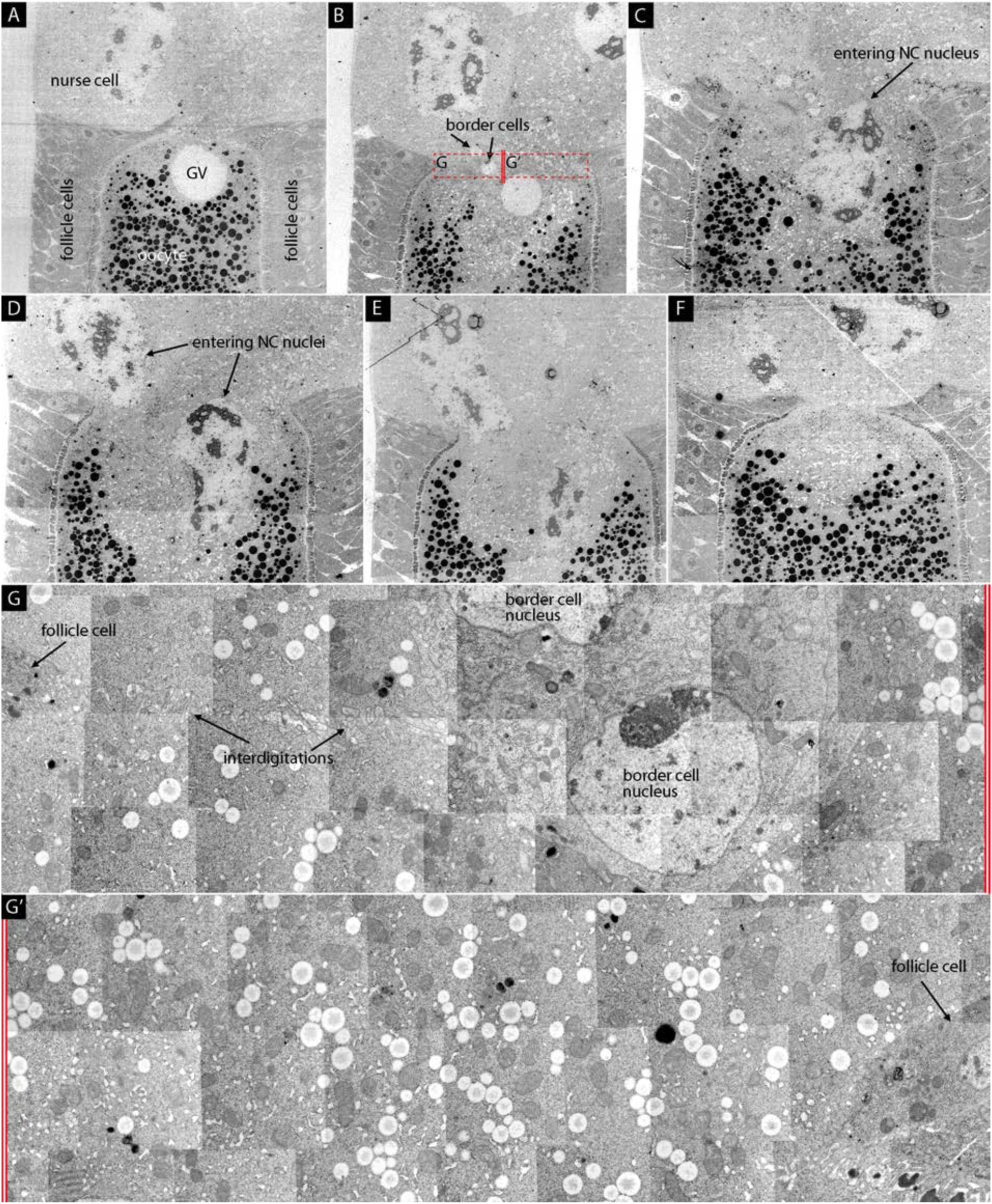
Fine structure images of oocyte with entering nurse cell nuclei. (A-F) Six successive frontal sections of a stage 10B egg chamber imaged by electron microscopy showing contiguous plasma membrane across the anterior oocyte face (A,) and nurse cell nuclei in open channel (B-F). (G,G’) Montage of higher magnification images from region indicated by dashed red lined box in (B) showing interdigitated membrane fragments and border cell nuclei on one side (G) and membrane-free ooplasm on the other (G’).

**Figure 8.**
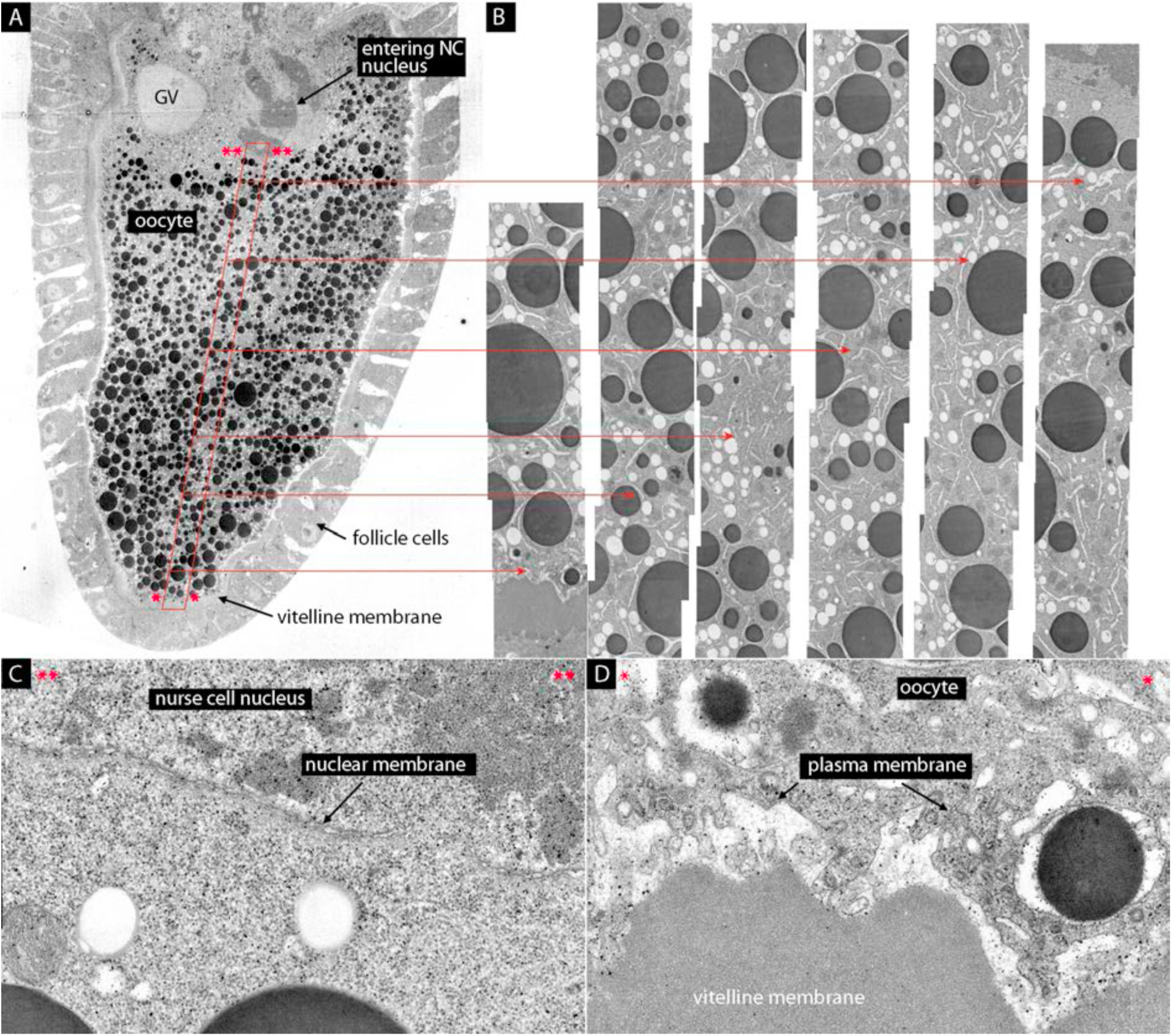
Fine structure images of ooplasm. Electron micrographs of a stage 10B egg chamber with an entering nurse cell nucleus showing the oocyte at low magnification (A) and a montage of high magnification images (B) showing the ooplasm devoid of membrane between the entering nurse cell nucleus at the anterior and plasma oocyte membrane juxtaposed to vitelline membrane at the posterior pole. Fine structure of the nuclear membrane of the entering nurse cell (C) and oocyte plasma membrane (D).

To eliminate the possibility that the plasma membranes that segregate the oocyte from the nurse cells in this region at earlier stages are either displaced posteriorly at stage 10B, a central column of the entire length of a stage 10B oocyte was imaged at high magnification and a montage was assembled (Fig. 8A,B). An entering nurse cell nucleus is visible at the most anterior end of the column, and its oocyte is visible at the interface with vitelline membrane material that is interposed between the oocyte and follicle cells (Fig. 8D). No plasma membrane is visible in the images of the ooplasm between the nurse cell nucleus at the anterior end and the posterior end of the oocyte (Fig. 8B). We conclude that the plasma membrane of the oocyte is not continuous across its anterior face and that there is an opening in the anterior face of the oocyte through which adjacent nurse cells enter.

The presence of a channel at the anterior face of the oocyte requires a transition between the plasma membranes of the oocyte and nurse cells. To examine the nature of this transition and to determine if the oocyte and nurse cells have separate and independent openings, or if the oocyte and nurse cell membranes fuse, we analyzed the edge of the channel in high resolution images such as those shown in Figure 9. A nurse cell nucleus extends posteriorly into the oocyte in the space that has follicle cells on both sides (Fig. 9A). Our interpretation of the membrane profiles in the region bounded by the red dashed lines is shown in Figure 9B: (1) the blue lines trace deep indentations of the nurse cell nuclear membrane, including cross sections through portions of the convoluted nuclear surface that appear to be detached in this section; (2) the green lines trace the plasma membranes of follicle cells on the left side of the channel; and (3) the red line traces the plasma membranes of the oocyte and nurse cell. As shown in the Figure 9C and 9D high magnification images the plasma membranes of the oocyte and nurse cell are continuous. The oocyte plasma membrane extends around the dense material of the vitelline membrane at the edge of the channel and around the finger-like protrusion of a follicle cell, connecting directly with the plasma membrane of the nurse cell that extends anteriorly and is juxtaposed with follicle cell plasma membranes. These images are consistent with the idea that the plasma membranes of the oocyte and nurse cells fuse to create a channel that directly connects oocyte and nurse cell cytoplasm.

**Figure 9.**
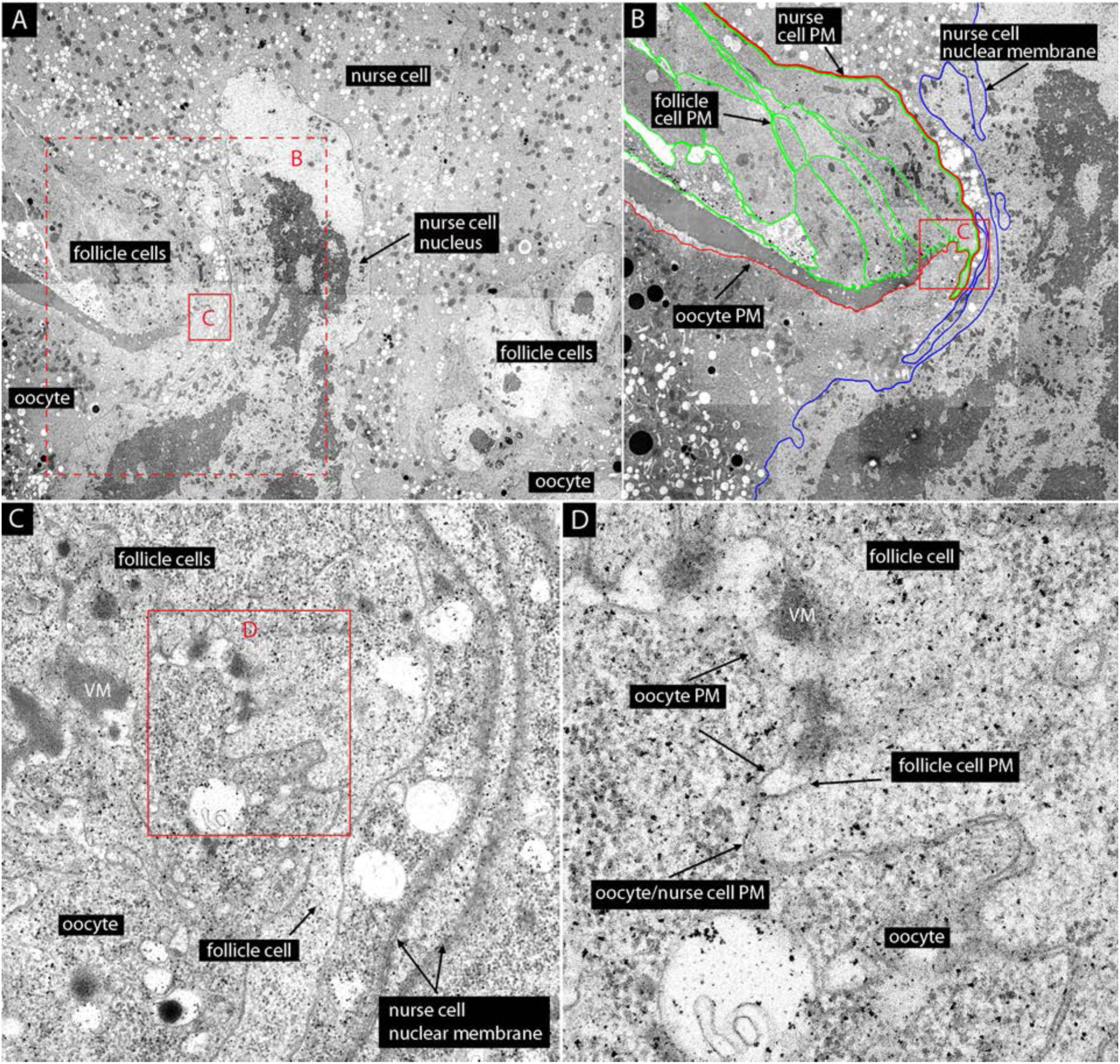
Contiguous plasma membrane of the oocyte and nurse cell. (A) An electron micrograph of an entering nurse cell nucleus. (B) Dashed red lined-box in (A) imaged at higher magnification with traced plasma membranes of follicle cells (green), contiguous plasma membranes of oocyte and nurse cell (red), and nuclear membrane of entering nurse cell (blue). (C) Red-lined box in (B) imaged at higher magnification. (D) Red-lined box in (C) imaged at higher magnification.

### Structure and fate of the entering nurse cell nuclei

We characterized the constitution of the entering nurse cell nuclei which change shape in stage 10B egg chambers (Figs. 1,3,6-9). Entering nuclei in the oocyte:nurse cell channel are elongated, with a dumbbell-like shape (Fig. 1E, 10A-A’’’). Although actin cables are prominent in other stage 10B nurse cell nuclei (Huelsmann et al., 2013), phalloidin staining did not detect actin cables in or around the entering nurse cell nuclei (Fig. 10A’’), and the uniform distributions of nuclear DNA and GFP are consistent with the idea that nuclear contents are not reorganized during extrusion through the nurse cell-oocyte channel. Staining for nuclear membrane markers such as Msp300 and lamin (Fig. 10B,B’), and fine structure analysis (Fig. 10C,C’) similarly reveals no apparent change to the nuclear membrane of the entering nuclei. However, the mAb414, which recognizes four different nucleoporin proteins of the nuclear pore complex (Davis and Blobel, 1986; Meier et al., 1995) and stains nurse cell nuclear membranes, does not stain the portion of the nurse cell nuclear membrane that is inside the oocyte (Fig. 10A’’’). This indicates that the molecular constitution of the nuclear envelope changes upon entering the oocyte.

**Figure 10.**
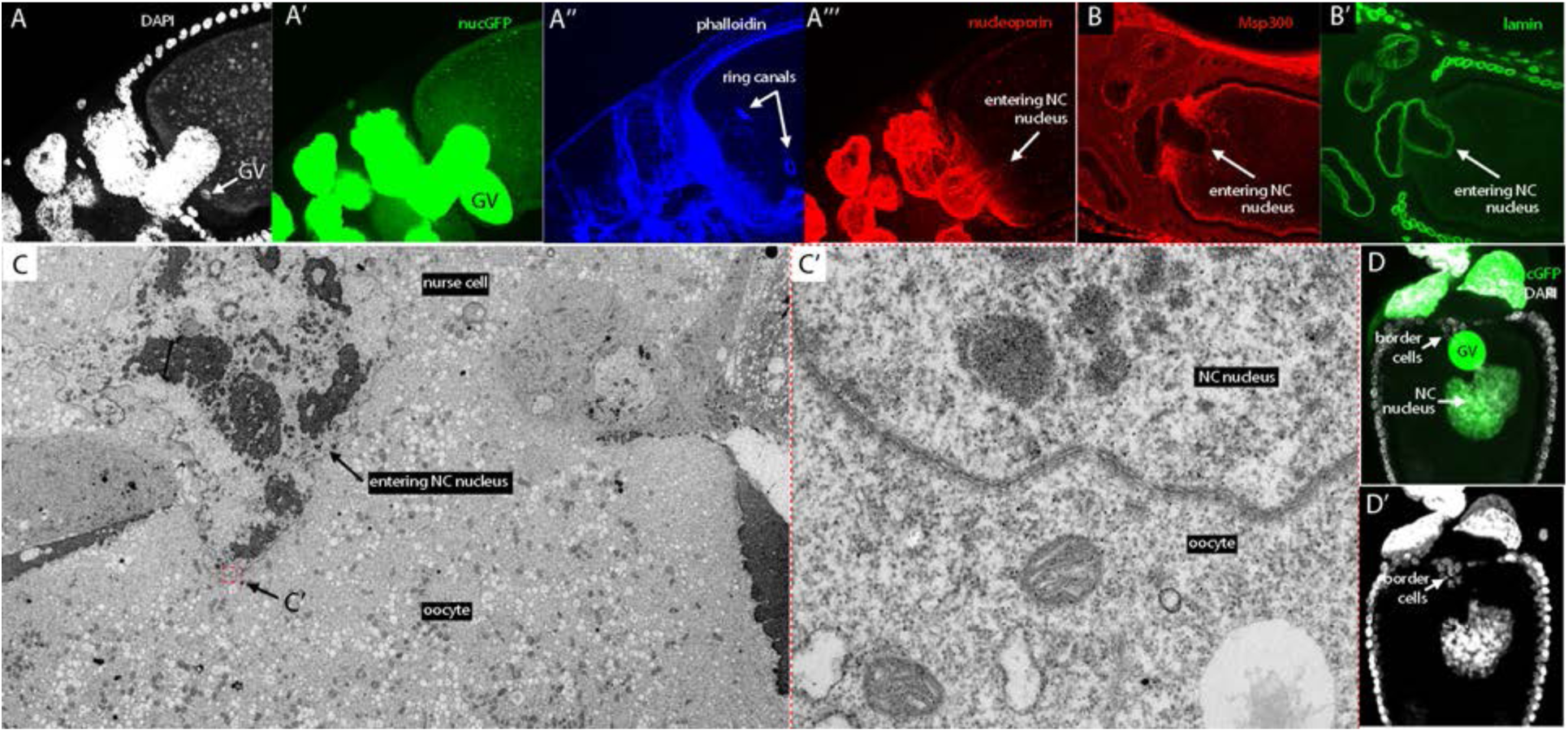
Anatomy of the nuclear membrane of the entering nurse cell nucleus. (A-A’’’) The entering nurse cell nucleus of the stage 10B oocyte is imaged with DAPI (white, A), nuclear GFP (green, A’), phalloidin (blue, A’’) and α-nucleoporin, which stains only the portion of the nucleus that has not entered (A’’’). (B,B’) Entering nurse cell nucleus stained with α-Msp300 (red) and α-lamin (green) antibodies. (C) Electron micrograph of entering nurse cell nucleus with red-lined region shown at high magnification in (C’). (D,D’) Nuclear GFP (green) and DAPI (white) marks nurse cell nucleus in the ooplasm, border cells, and GV.

Nurse cell nuclei that are deep in the oocyte interior are not round spheres and are larger than other nurse cell nuclei (Figs. 1,3), and fluorescence of DAPI and nuclear GFP is low and diffuse relative to nuclei in nucleated nurse cells (Fig. 10D,D’). The nurse cell nuclei in the oocyte dissipate, but we were not able to identify how they are eliminated. Studies of nurse cell elimination have not found the process to be either apoptotic or autophagic (Peterson and McCall, 2013), and we did not observe that the entering nurse cell nuclei stain with anti-caspase antibody.

### The oocyte:nurse cell interface of stage 11 egg chambers

As noted above, immunofluorescence images of stage 11 egg chambers are consistent with the idea that the oocyte:nurse cell channel shrinks after stage 10B, and that the membranes of the oocyte and nurse cells close the channel of stage 10B egg chambers (Figs. 4G, 5D). The ring canals that precede the nurse cell nuclei into the oocyte also appear to return to their original positions and to reform the patterned quartet of ring canals at the interface (Fig. 5G). However, because these images do not resolve the resolution the separate plasma membranes of the oocyte and nurse cells, they do not show whether the channel is completely sealed. We examined EM sections of stage 11 egg chambers, and found that although the stage 11 egg chamber does not have a large channel at the oocyte:nurse cell interface, and although the vitelline membrane (VM) and follicle cells extend across the anterior face of the oocyte, a small gap remains (Fig. 11A-E). These images are consistent with the idea that the channel does not completely close and that the ooplasm is not completely separated from nurse cell cytoplasm by plasma membranes. The simplest interpretation is that there is continuity between ooplasm and enucleated nurse cells 2 and 5.

**Figure 11.**
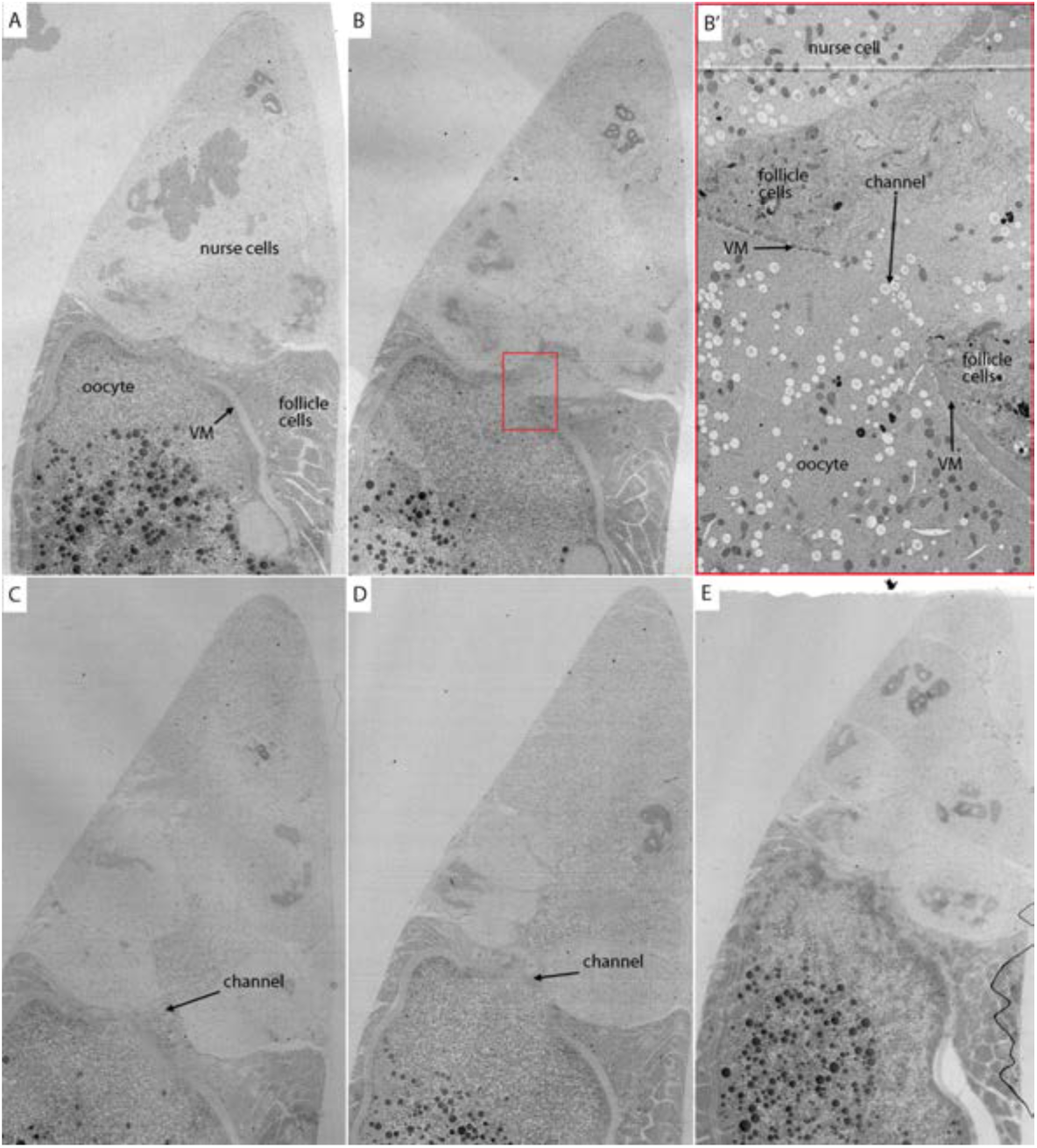
Fine structure of anterior face of a stage 11 oocyte. (A-E) Successive sections imaged by electron microscopy of a stage 11 egg chamber showing contiguous plasma membrane at the anterior oocyte face in outer sections (A,E) and channel in intermediate sections (B-D). (B’) Higher magnification image of red-lined boxed region in (B) showing nurse cell, follicle cells, vitelline membrane (VM), oocyte, and channel.

### Nurse cell nuclear elimination is essential for oocyte maturation

Counts of stage 11 egg chambers in Figure 3G show that approximately 90% had less that 15 nurse cell nuclei, and 10% did not reduce the number nurse cell nuclei. We analyzed stage 11-14 egg chambers in WT ovaries to determine whether the process of nuclear elimination is necessary for egg chamber development. The total number of nurse cell nuclei can be determined in DAPI-stained, fixed preparations of stage 11-13 egg chambers that also express nuclear-localized GFP. Figure 12A shows a typical stage 12 egg chamber that has 13 nurse cell nuclei. The egg chamber shown in Figure 12B is not normal: it is classified as stage 12-13 because of the arrangement of the follicle cells around the incipient dorsal appendages, but it has 15 nurse cell nuclei that are not arranged normally in the usual ordered, compact organization. In addition, there has been no expansion of the oocyte or compensatory reduction of the nurse cell compartment that is characteristic of normal egg chambers. WT ovaries also have stage 14 oocytes that are similarly truncated and have abnormal dorsal appendages (Fig. 12 C,D), and the position of the DAPI-stained oocyte nucleus is abnormal in these oocytes (Fig. 12 E,F). The fraction of abnormal egg chambers at stages 11-13 and stage 14 increased as females aged (Fig. 12G).

**Figure 12.**
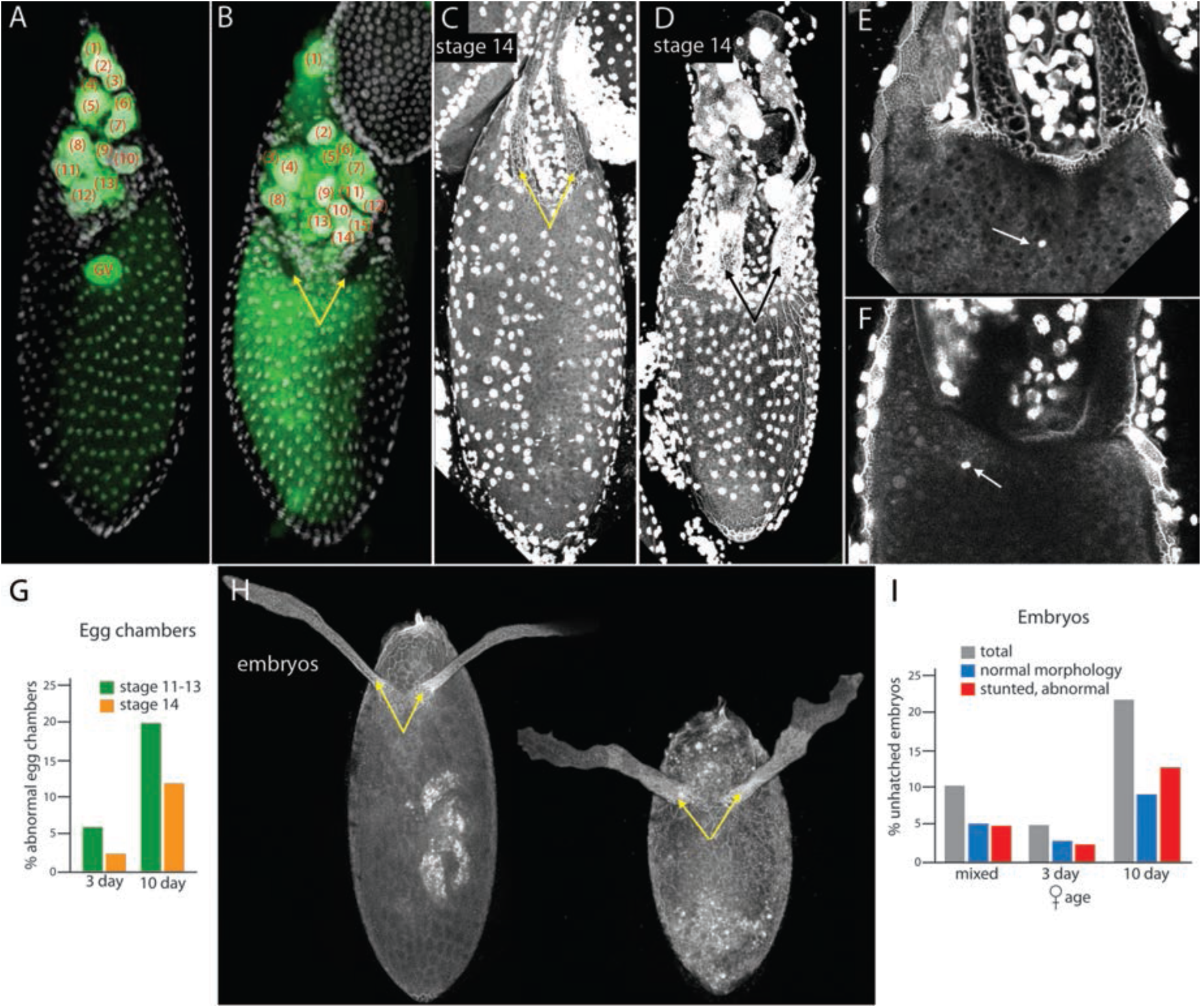
Nurse cell numbers vary in WT egg chambers. (A,B) Stage 12 egg chambers stained with nuclear GFP (green) and DAPI (white) with counts (parentheses) showing 13 nurse cell nuclei in (A) and 15 in (B) that has abnormally short oocyte and abnormal morphology of regions forming dorsal appendages (arrows). (C,D) Stage 14 oocytes with normal morphology (C) and abnormal morphology (D); arrows indicate dorsal appendages. (E,F) High magnification images of anterior regions of normal (E) and abnormal (F) stage 14 oocytes with oocyte nucleus (arrows) position normally in the center of (E) and not centered in (F). (G) Graph indicating percentage of egg chambers with abnormal morphology in ovaries of 3 day and 10 day old females. (H) Embryos with normal (left) and abnormal (right) morphology manifested in shape of dorsal appendages (arrows) and size. (I) Graph tabulating percentage and morphology of unhatched embryos laid by females of mixed age, and 3 and 10 days posteclosion.

To determine the fate of the abnormal stage 14 oocytes, embryos deposited by WT females were collected and analyzed. Approximately 90% of the embryos from actively laying females of indeterminate (mixed) age hatched and developed to the 1^st^ instar larval stage, 5% developed but did not complete embryogenesis, and 5% were undersized, had abnormal dorsal appendages, and were not apparently fertilized (Fig. 12H,I). Young females (3 days post eclosion) laid fewer unhatched and abnormal eggs than did older females (10 days post eclosion). Due to the stunted appearance, abnormal dorsal appendages, and frequency, we propose that the abnormal-shaped unhatched eggs are the products of the stage 11-13 egg chambers with 15 nucleated nurse cells and the abnormal stage 14 oocytes.

## Discussion

The mature Drosophila egg is the product of a multi-day process to which several types of germ line cells contribute. The separate fates of the oocyte and sacrificial nurse cells are set at the first division of the primary oogonial cell (de Cuevas and Spradling, 1998), but our finding that specific nurse cells undergo enucleation at stage 10B is the first evidence that at least these nurse cells follow a program that is different from the others; the infertility of oocytes that do not execute the enucleation program is evidence of its importance. We can only speculate at this time on its purpose.

Although the specific nurse cell nuclei dissipate upon entering the oocyte, we do not consider it likely that they enter the oocyte to contribute their components, as these nurse cells are each connected to the oocyte by a ring canal that provides a direct conduit for its products. It also seems unlikely that the nuclei are eliminated in order prevent them from making something deleterious. We consider a third possible reason – that enucleation at stage 10B is part of a process that creates an efficient outlet for rapid transfer of materials from the fifteen nurse cells to the oocyte.

Stage 10B is a transition when the growth rate of the oocyte increases 2x from a doubling time of approximately 5.5 hours that does not vary for stages 2 to 10A (Fig. 5E). The doubling time of oocytes at 10B and subsequent stages that we measured is approximately 2.8 hours, a rate that is less than the 30 minutes reported previously (Guild et al., 1997), but that is nonetheless a significant increase over the previous stages. The change in oocyte growth rate contrasts with the relatively constant rate of increase in egg chamber volume between stages 2 and 14. This difference is consistent with the idea that stage 10B initiates the dumping period of rapid transport of nurse cell components into the oocyte (Matova and Cooley, 2001b).

Dumping has been attributed to concerted “contraction” and ‘explusion” through ring canals that increase gradually in size throughout the earlier stages (Warn et al., 1985). However, the ring canals of stage 10B egg chambers are approximately 10 μm diameter, much smaller than the approximately 30 ⍰m diameter nurse cell nuclei that enter the oocyte. We found that the nurse cell nuclei move into the oocyte through a large channel that directly connects nurse cell cytoplasm with ooplasm, that they do not move into the oocyte through ring canals (Figs. 6-10). The channel opens coincident with the initiation of dumping, and although we do not know if it is present later in stage 12 and stage 13 egg chambers, we found that it is present in stage 11 egg chambers. The ring canals persist during these stages and presumably continue to provide a means for cytoplasm transfer, but the channel could be a second type of portal during the dumping period.

This potential role for the channel does not suggest why specific nurse cells become enucleated, but it is consistent with the phenotype of oocytes in egg chambers that do not reduce the number of nurse cell nuclei. Stage 12 and 13 egg chambers that retain 15 nurse cell nuclei are similar in size to stage 12-13 egg chambers with 13 nurse cell nuclei, but the oocytes in egg chambers with 15 nurse cell nuclei are shorter and the nurse cell compartment is proportionally elongated (Fig. 12). The stunted shape of these oocytes is consistent with the idea that the dumping phase of rapid cytoplasmic transfer failed, and with the possibility that nurse cell nuclei did not enter these oocyte because the channel did not to form. It may be relevant that the number of nurse cell nuclei in stage 11 egg chambers varies: ∼60% have 13, ∼20% have 14, and ∼6% have 12, and the fraction of egg chambers that have 15 nucleated nurse cells is the same as the fraction of oocytes that are stunted. We suggest that oocyte maturation may proceed normally in egg chambers with 12, 13, and 14 nucleated nurse cells, but not in egg chambers that do not successfully execute the nuclear extrusion process.

If this rationale for enucleation seems prosaic, the biology that enables it is fascinating and complex. For example, the cell fusion that joins the stage 10B oocyte with nurse cells is unique in several respects. First, it is pre-patterned by the arrangement of the specified nurse cells that fuse with the oocyte. Second, its scale is massive due to the dimensions of the channel that joins the large oocyte and large nurse cells. Third, it appears to be reversible in the sense that the enucleated nurse cells of stage 11 egg chambers have shapes and appearances that are similar to their nucleated neighbors, despite the small opening that remains. Finally, the fusion-generated channel not only makes nuclear extrusion and cytoplasmic mixing possible, but also brings the cluster of border cells, which are of somatic origin, and the germinal vesicle of the oocyte into close proximity (Fig. 6A-F). It is known both that the border cells move in a stereotyped sequence when they reach the anterior face of the oocyte and that failure to complete the sequence does not allow for normal oocyte maturation (Duchek and Rørth, 2001; Montell, 2003). However, the position of the border cells in the channel is a novel feature that raises the possibility of a role for the border cells in the enucleation program, or alternatively, of a role for the cell fusion/enucleation program in bringing the germinal vesicle and border cells together. Subsequent work in our lab (Ali-Murthy, Fetter, and Kornberg, in preparation) supports the latter model, but we mention these speculations here to articulate some of the many new types of questions these findings raise.

## Materials and Methods

### Reagents

#### Fly lines

MTD-Gal4 (Bloomington #31777), UASp-NLSGFP (from J. Vasquez), UASp-Lifeact/TM6 (from W. Sullivan), UAS-CD8:GFP (Roy et al., 2011), Act5C-Gal4/CyO, Dpp:Cherry (Fereres et al., 2019)

#### Antibodies

mouse α-lamin, mouse α-HTS, and rat α-cadherin (Developmental Studies Hybridoma Bank); α-MSP300 (Y. Gruenbaum), mouse mAb 414 (M. D’Angelo)

### Histology

#### Antibody staining

Five to ten pairs of ovaries were dissected in PBS together with their common duct, the linked ovaries were transferred with forceps to a watch glass with 16% formaldehyde, and after 5 minutes, the ovarioles were gently separated with fine forceps, transferred to an Eppendorf tube and rotated slowly for 30 minutes, followed by at least three 10 minutes washes in PBT and processing as described (Ali-Murthy et al., 2013). All images were obtained with a Leica SPE confocal microscope.

#### Ovary preparation

“Normal” preparations were from flies 4-6 days post eclosion that had been incubated at 25°C in bottles (20-25 females and 15-20 males) and transferred to bottles with fresh medium 2-3 days prior to dissection. “Optimized” preparations were from flies (20-25 females and 15-20 males) 3 days post eclosion that had been transferred to bottles with fresh food.

Full ovary preparations (as in Figure 1A) were dissected in PBS, several paired ovaries (linked by their common duct) were transferred with forceps to an Eppendorf tube with 16% formaldehyde for 30 minutes, washed 3×10 minutes with PBT, stained with DAPI (10 minutes), washed 3×10 minutes with PBT, and transferred by pipet to a microscope slide (one pair/slide). Liquid surrounding the ovary pair was removed with Kimwipe tissue and replaced with Vectashield mounting medium. Support, either a cover slip or layer of tape, was placed to either side of the ovaries, followed by a standard 1.5H cover slip and sealing with nail polish.

To view unfixed (live) preparations, ovaries were dissected in PBS directly on a microscope slide, the outer sheath was disrupted with either a tungsten needle or fine forceps to free the ovarioles, supports were placed to either side, and a cover slip placed over the preparation.

To analyze nurse cell nuclei, fixed ovariole preparations from flies that express nuclear-localized GFP (MTD-Gal4>UASp-NLSGFP) were stained with antibody and mounted under a supported coverslip for imaging. Approximately 250 confocal optical sections separated by 0.27 ⍰m were recorded, and the images were assembled with ImageJ to create a 3D rendering. To count nurse cell nuclei, the 3D structure was rotated to identify each nucleus in preparations that were stained with phalloidin to mark cell borders and the number of ring canals associated with each nucleus was determined. Each of the 16 germline cells has a unique signature composed of the number of ring canals and nearest neighbors with 1, 2, 3, or 4 ring canals.

Egg chamber volumes were determined by measuring the total length (L_OC_), width (W), and depth (D) of DAPI-stained preparations, and calculated by assuming an ellipsoid shape (4/3 × π × L_OC_ × D × W). Oocyte volumes were determined by measuring the length of the oocyte (L_O_) and calculating the volume as 4/3 × π × L_O_ × D × W. For each stage, n≥10.

To analyze egg chamber phenotypes, WT females 3 and 10 days posteclosion that had been transferred daily to bottles with fresh medium were stained with DAPI and imaged by confocal microscopy. 40 ovaries were analyzed for each time point. Egg chambers analyzed: 3 day: 96 stage 11-13, 104 stage 14; 10 day: 192 stage 11-13, 152 stage 14. To analyze embryo phenotypes, 100 embryos from 4 hour collections laid by WT females 3 days, 10 days, and <14 days (mixed) posteclosion were incubated for 18 hours. Unhatched embryos were mounted directly in PBS on a microscope slide and imaged under a supported coverslip. Total unhatched embryos: mixed: 36/300 embryos; 3 day: 11/200 embryos; 10 day: 66/300 embryos

### Electron microscopy

Ovaries were removed from WT females under a dissecting microscope in PBS on a microscope slide set on an ice-chilled plate. The slide was transferred to a transmitted light dissecting microscope, the ovary was gently disrupted to slightly separate ovarioles, stage 10B egg chambers were identified, and the stage 10B egg chambers with oocytes that had slightly cleared, small regions near their anterior face were transferred with a cut-off Eppendorf pipet tip in a volume of 5 μl to immersion fixation consisting of either 2% glutaraldehyde in 0.1 M Na-cacodylate buffer, pH 7.3 for 1 hour at room temperature (RT) or 1% glutaraldehyde/1% OsO_4_ in 0.1 M Na-cacodylate buffer, pH 7.3 for 45 minutes on ice. After fixation, samples were rinsed in 0.1 M Na-cacodylate buffer, followed by post-fixation in 1% OsO_4_ in 0.1 M Na-cacodylate buffer for 1 hour at RT. The samples were then rinsed with water, *en bloc* stained with 5% uranyl acetate in water for 1 hour at RT. Following *en bloc* staining, the samples were rinsed with water, dehydrated in an ethanol series followed by propylene oxide, infiltrated and then polymerized in Eponate 12 resin (Ted Pella, Inc., Redding, CA). 50 nm sections were cut with a Leica UCT ultramicrotome (Leica Microsystems, Buffalo Grove, IL) using a Diatome diamond knife (Diatome USA, Hatfield, PA), picked up on Pioloform coated slot grids and stained with uranyl acetate and Sato’s lead (Sato, 1968). Sections were imaged with a FEI Tecnai 12 TEM at 120 kV using a Gatan 4k × 4k camera. TrakEM2 in Fiji was used to align image montages (Cardona et al., 2012; Schindelin et al., 2012).

## Acknowledgements

We thank: W. Sullivan, S. Younger, and the Bloomington Stock Center for fly stocks; M. D’Angelo mAb 414 antibody, Y. Gruenbaum for α-Msp300 antibody and the Developmental Studies Hybridoma Bank for α-lamin, α-HTS, and α-cadherin antibodies. This work was funded NIH grants R01GM109410 and R35GM122548 to T.B.K..

## Competing interests

The authors do not have any financial or non-financial competing interests.

---

**Figure 2 supplement.**
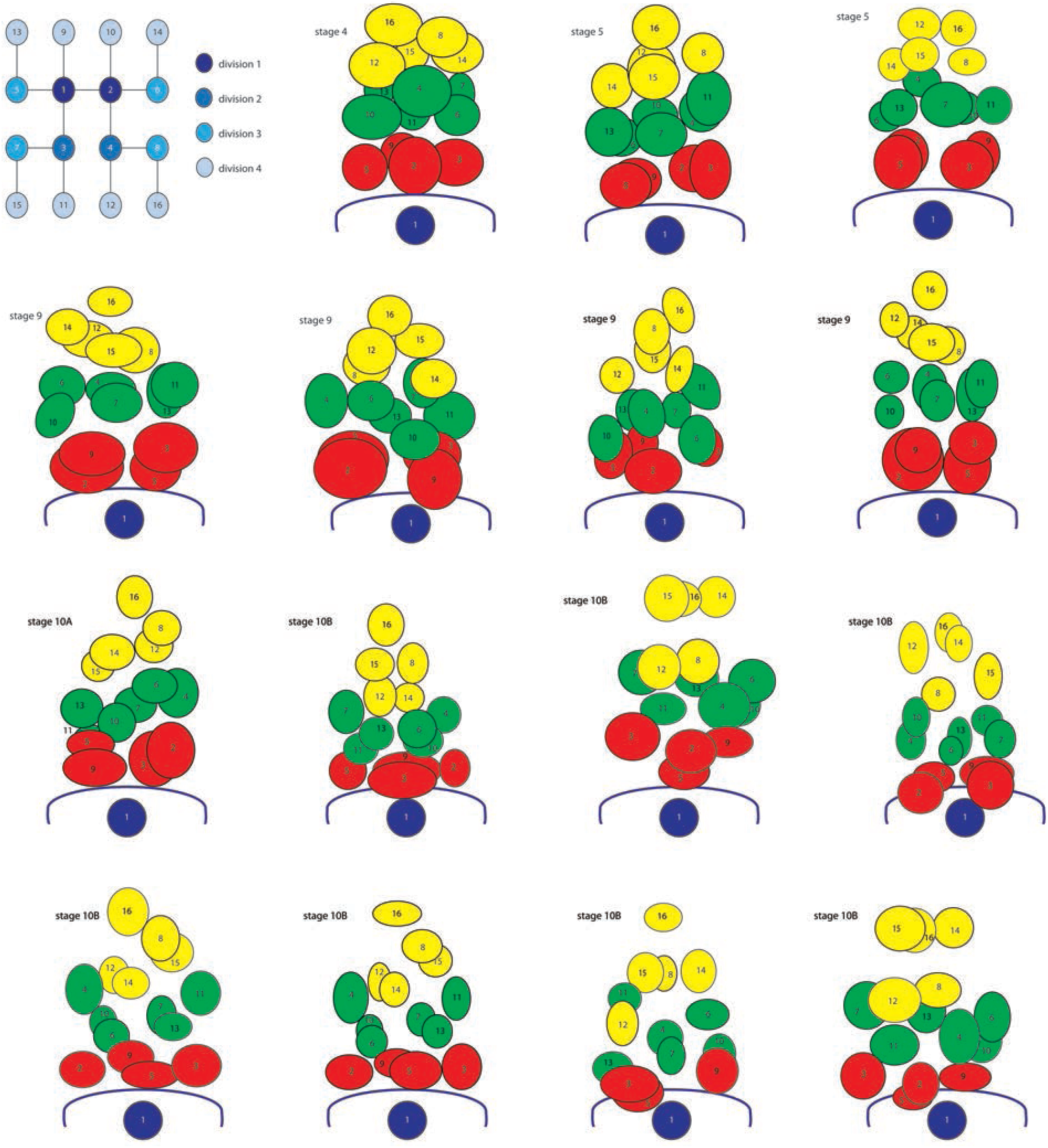
Positions of nurse cell nuclei stages 4-10B. Upper left: germline lineage tree. Cartoon depictions of stage 4 – stage 10B egg chambers.

